# Convergent evolution of skim feeding in baleen whales

**DOI:** 10.1101/2022.08.16.504064

**Authors:** Ludovic Dutoit, Kieren J. Mitchell, Nicolas Dussex, Catherine M. Kemper, Petter Larsson, Love Dalén, Nicolas J. Rawlence, Felix G. Marx

## Abstract

The origin of pygmy right whales (Caperea marginata), the smallest and most enigmatic of the living baleen whales, remains contentious. Morphological analyses largely continue to ally Caperea with right whales (balaenids) based on shared cranial features like a tall braincase and a narrow, highly arched rostrum. By contrast, molecular data and some anatomical evidence suggest a closer relationship with rorquals (balaenopterids), but fail to explain “the substantial issue of convergence posed by the many balaenid features of Caperea” (Berta & Demere, 2017). To resolve this question, we sequenced the nuclear genome of C. marginata (812,269,251 paired reads; 47X average depth of coverage, with 89.33% of the genome covered at ≥10X) and subjected it to a multispecies coalescent analysis including representatives of all baleen whale families. Our results confirm Caperea as sister to rorquals and, thus, the convergent origin of its right whale-like anatomy. Considering this overwhelming molecular evidence, we propose that the traditional taxonomic grouping of Caperea with right whales be abandoned.

---

The origin of pygmy right whales (*Caperea marginata*), the smallest and most enigmatic of the living baleen whales, remains contentious. Morphological analyses largely continue to ally *Caperea* with right whales (balaenids) based on shared cranial features like a tall braincase and a narrow, highly arched rostrum (Bisconti et al., 2017). By contrast, molecular data and some anatomical evidence suggest a closer relationship with rorquals (balaenopterids) (McGowen et al., 2020, Marx & Fordyce, 2016), but fail to explain “the substantial issue of convergence posed by the many balaenid features of *Caperea*” (Berta & Deméré, 2017). To resolve this question, we sequenced the nuclear genome of *C. marginata* (812,269,251 paired reads; 47X average depth of coverage, with 89.33% of the genome covered at ≥10X) and subjected it to a multispecies coalescent analysis including representatives of all baleen whale families (Árnason et al., 2018; see Supporting Information for full details). Our results confirm *Caperea* as sister to rorquals (Fig. 1A) and, thus, the convergent origin of its right whale-like anatomy. Considering this overwhelming molecular evidence, we propose that the traditional taxonomic grouping of *Caperea* with right whales be abandoned.

**Figure 1.**
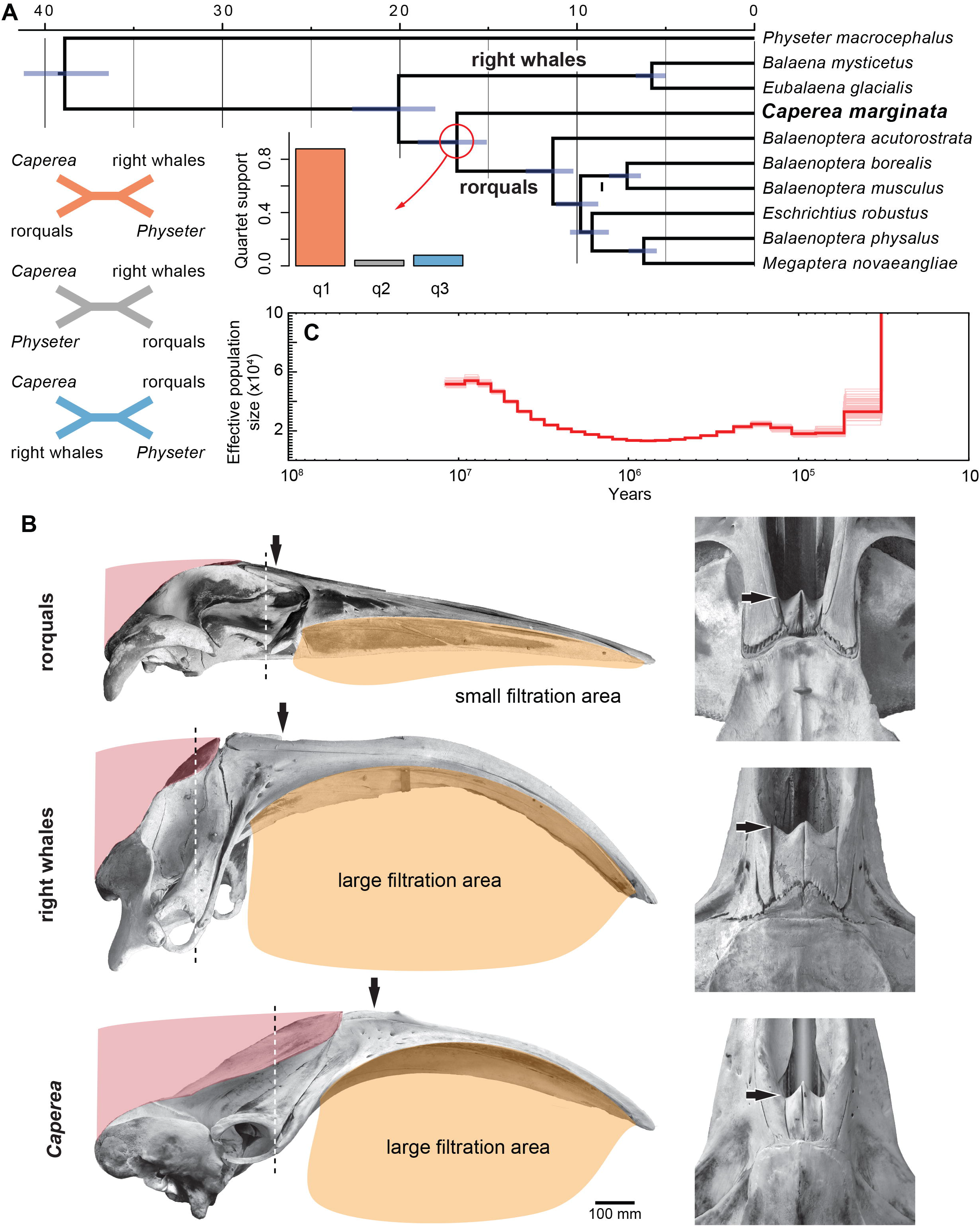
Origin and demographic history of pygmy right whales. (A) Species tree of baleen whales based on 89,115 individual gene trees depicted with node ages estimated using exons from 344 protein-coding genes that exhibited clock-like evolution in the sampled taxa. Blue bars represent 95% highest posterior densities. Quartet scores for all possible arrangements of *Caperea*, rorquals, right whales, and the sperm whale (*Physeter macrocephalus*) are shown to the left of the tree—q1 is the arrangement depicted in the species tree. (B) Comparative skull architecture of rorquals (top), right whales (centre), and *Caperea* (bottom), showing the relative sizes of the baleen apparatus/ filtration area (yellow) and the neck muscles (red). Black arrow denotes the posterior margin of the nares, the dashed line the anterior margin of the orbit. Note how the neck muscles extend anterior to the level of the orbit in right whales and *Caperea*, and how the nares in both are shifted far forward as a result. (C) Ancestral effective population size of *Caperea.* The dramatic increase in estimated effective population size at ~30 ka is likely an artefact resulting from a lack of recombination and coalescence events in the recent past.

The resemblance between *Caperea* and balaenids likely arises from their similar skim feeding strategies, which require a large cross-flow filtration surface (Werth & Potvin, 2016). In both cases, this is achieved via notably elongate baleen plates, whose size and functional requirements shape the remainder of the skull. Thus, the large baleen rack is accommodated by an arched rostrum, which in turn requires a tall braincase and anteriorly positioned neck muscles for support. Further knock-on effects are evident on the skull vertex, with the blowhole and surrounding rostral bones being ‘pushed’ forward by the neck muscles and, thus, anterior to the eyes (Fig. 1B). We suggest that all these traits are interrelated and ultimately driven by the enlargement of the baleen apparatus for skim feeding, yet have frequently been treated as independent phylogenetic characters apparently uniting *Caperea* with balaenids.

Our results call for a re-evaluation of morphology-based cladistic datasets to control for the effects of convergence and uncover the true origin of pygmy right whales in the fossil record. We estimate that *Caperea* diverged from rorquals between 19–16 Ma (Fig. 1A; Supporting Information), yet the oldest pygmy right whale fossils are no older than 8–7 Ma or, perhaps, 10–9 Ma (Tsai et al., 2017). This long ghost lineage may either reflect genuine undersampling of the fossil record or suggest that the earliest members of the *Caperea* lineage have been misidentified, as previously suggested with regard to cetotheriids (Marx & Fordyce, 2016). We cannot evaluate these hypotheses directly, but note that both the phylogenetic position of *Caperea* and its estimated time of divergence are consistent with a cetotheriid origin, with the oldest members of that family dating to the Middle Miocene (ca 14.8–12.5 Ma) (Collareta et al., 2021, Gol’din, 2018).

In addition to resolving the taxonomic affinities of *Caperea*, our new genomic data allow us a first insight into its demographic history. Our estimates for the trajectory of *Caperea* effective population size (*N_e_*) over the last 10 Ma resemble those of fin, minke, and sei whales (Árnason et al., 2018)—a gradual decline in N_e_ since the Late Miocene plateaus at around 800 ka, followed by an increase lasting until around 200 ka (Fig. 1C). These fluctuations in *N*_e_ may reflect changes in census population size driven by past climate change and/or changing rates of gene flow between different populations (Mazet et al., 2016). While *Caperea* is presently distributed exclusively in the Southern Ocean, where there are presumably few barriers to gene flow, the discovery of Pleistocene *Caperea* fossils from Italy and Japan (Tsai et al., 2017) hints at past population structure. Periodic cooling during Pleistocene glacial cycles may have created opportunities for cold-adapted species to cross the equator, resulting in complex population dynamics for otherwise temperate to polar species like *Caperea marginata*.

## ACKNOWLEDGEMENTS

D. Stemmer, S. South, K. Roberts and J. Sumner helped to collect samples. The National Genomics Infrastructure in Stockholm (funded by Science for Life Laboratory, the Knut and Alice Wallenberg Foundation and the Swedish Research Council) and SNIC/Uppsala Multidisciplinary Center for Advanced Computational Science assisted with massively parallel sequencing and access to the UPPMAX computational infrastructure.

## AUTHOR CONTRIBUTIONS

L.Du. and K.J.M. conducted the phylogenetic analyses; N.D. and L.Da. performed bioinformatics data processing and conducted the demographic analysis; C.K. provided samples and input on the biology of *Caperea*; P.L. performed DNA extraction and library preparation; F.G.M. and N.J.R conceived of the study and organised the collaborative project. All authors discussed and wrote the paper.

## SUPPORTING INFORMATION

Additional supporting information may be found online in the Supporting Information section at the end of this article.

## Notes

### Competing Interest Statement

The authors have declared no competing interest.

