## Supplementary Material for "Convergent evolution of skim feeding in baleen whales"

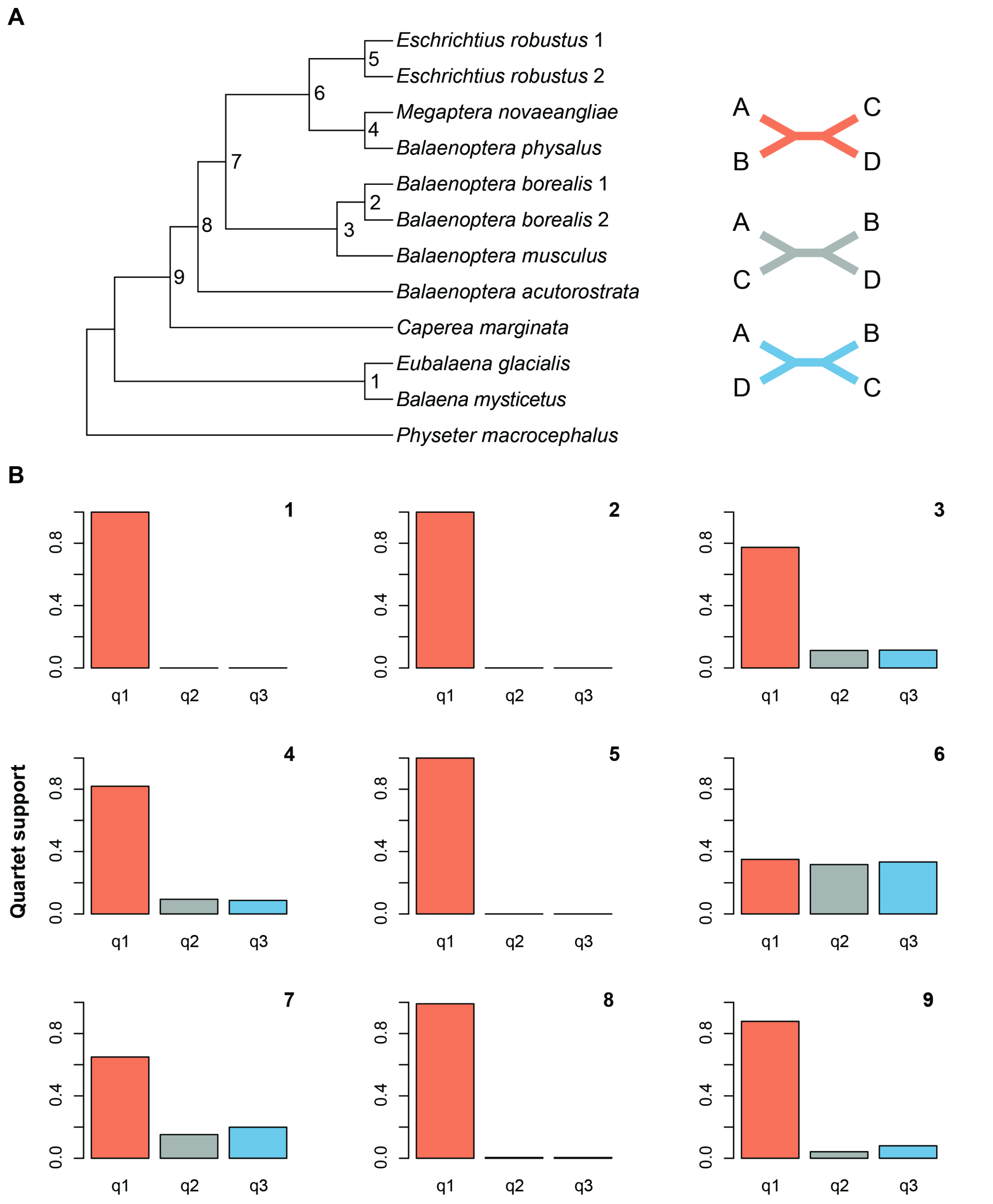


**Figure S1. Origin and demographic history of pygmy right whales.** (A) MSC species tree and (B) ASTRAL quartet support scores.

### Material and Methods

*Sample collection, DNA extraction, library preparation and high-throughput sequencing*

We extracted genomic DNA from kidney tissue obtained from a juvenile female pygmy right whale (South Australian Museum, Adelaide, specimen M27462) that came ashore near Port Vincent, South Australia, in September 2017. The extraction was performed using a Kingfisher robot (Thermo Fisher Scientific), following the manufacturer's Kingfisher blood & tissue extraction protocol. Double-stranded Illumina libraries were built using the Illumina TruSeq PCR-free library preparation protocol and then sequenced on an Illumina HiSeqX in High Output mode with 2 × 150 bp paired-end sequencing chemistry at Sci*Life*Lab (Uppsala University, Sweden).

*Data mapping and consensus genomes generation*

Raw sequence data from the pygmy right whale were demultiplexed using bcl2Fastq v2.17.1 from the CASAVA software suite with default settings (Illumina Inc.). Mapping and variant calling were carried out in a beta version of the GenErode bioinformatics pipeline (Kutschera et al., 2022). We mapped genomic data of ten whale species (12 genomes; Table S1), including the pygmy right whale, to the assembly of the sperm whale (*Physeter macrocephalus*; GCF_002837175.2). First, we identified repeat regions in the reference assembly using RepeatModeler v1.0.11 and RepeatMasker v4.0.8 (Smit & Hubley, 2008-2015). Next, adapter trimming was done with trimmomatic v0.32 (Bolger et al., 2014), trimmed reads were mapped with BWA v0.7.17 (Li & Durbin, 2010) using the mem algorithm, and PCR duplicates were sorted and removed using SAMtools v1.12 (Li et al., 2009). Finally, reads were realigned around indels using GATK IndelRealigner v3.4.0 (McKenna et al., 2010).

Variants were called with the *bcftools mpileup* command of bcftools v1.13 (Li et al., 2009, Li, 2011), including orphaned reads. We used a minimum depth of coverage (DP4), roughly corresponding to ⅓ of the average depth of coverage, and filtered SNPs within 5 bp of indels. We used BEDTools v2.29.2 (Quinlan & Hall, 2010) *complement* tool to identify all sites missing from the VCF file, and then further used BEDTools to mask repeat sites based on the repeat mask of the reference sperm whale. Finally, we used *bcftools consensus* to generate consensus genomes for each of the 12 genomes from the resulting bam files and – following (Dussex et al., 2021) – masked polymorphic sites with 'N' to avoid the reference site being called for missing sites in any of the mapped genomes (https://github.com/ndussex/whale_genomics.git).

**Table S1.** Sources of genomic data. ENA, European Nucleotide Archive

| Species | ENA accession number | Study |
|---|---|---|
| *Balaena mysticetus* | SRR1685383 | http://www.bowhead-whale.org/ |
| *Balaenoptera acutorostrata* | SRR893003 | (Yim et al., 2013) |
| *Balaenoptera borealis* | SRR5665645, SRR5665646 | (Árnason et al., 2018) |
| *Balaenoptera musculus* | SRR5665644 | (Árnason et al., 2018) |
| *Balaenoptera physalus* | SRR5665643 | (Árnason et al., 2018) |
| *Caperea marginata* | PRJEB46892 | This study |
| *Eschrichtius robustus* | SRR5665641, SRR5665642 | (Árnason et al., 2018) |
| *Eubalaena glacialis* | SRR5665640 | (Árnason et al., 2018) |
| *Megaptera novaeangliae* | SRR5665639 | (Árnason et al., 2018) |
| *Physeter macrocephalus* | SRR6117283* | (Árnason et al., 2018) |

*reference assembly used for mapping: GCF_002837175.2

*Phylogeny*

The genomes were split into non-overlapping 20 kb windows. Windows for which any species had fewer than 8,000 called (i.e. non-N) sites were excluded, leaving 89,115 windows. Trees were created for each window using RAxML v8.2.12 (Stamatakis, 2014) with the GTR+G substitution model and rooted using the sperm whale as outgroup. The best-scoring tree for each 20 kb window was analyzed using ASTRAL v5.7.5. Quartet scores and trees were visualized using the *ape* v5.3 (Paradis & Schliep, 2019) package implemented in R v 3.5.2 (R Development Core Team, 2021).

*Molecular dating*

To estimate divergence times, we first extracted the exons for each protein-coding gene (n=20,196) from each consensus sequence based on their coordinates in the annotated sperm whale assembly. We then used a rapid bootstrap analysis in RAxML v8.2.0 (-f a –m GTRGAMMA -# 100) to create a phylogenetic tree for each gene. We subjected these trees to three filters using PhyKIT v1.11.5 (Steenwyk et al., 2021) and the adephylo v1.1-11 package (Jombart et al., 2010) in R v3.6.3: mean bootstrap support >80%, measured using the *bipartition_support_stats* command in PhyKIT to remove genes with low information content; Robinson-Foulds distance to the species tree ≤4, measured using the *robinson_foulds_distance* command in PhyKIT to remove genes that deviated excessively from the species tree; and a coefficient of variation of the root-to-tip distance <0.1, measured using the *dist.root* function in adephylo to remove "non-clocklike" genes with excessive rate heterogeneity.

The 344 genes that passed our filters were analyzed using *baseml* (part of the PAML v4.8a package; (Yang, 2007)) with the GTR substitution model and a global molecular clock to produce an estimate of per-gene rates of evolution. Genes were then ranked from fastest to slowest and divided into eight bins of 43 genes each with similar rates of evolution. The genes in each bin were concatenated for downstream analysis as eight separate partitions using *mcmctree* (part of PAML) and alignments were sub-sampled such that our final dataset only included a single representative of each species (individuals with the most missing data were removed).

**Table S2.** Divergence date estimates assuming autocorrelated vs independent rates and three alternative maximum root ages. Abbreviations: AC, autocorrelated; C, Cetacea; M, Mysticeti.

| | **Mean age (95% highest posterior density)** | | | | | |
|---|---|---|---|---|---|---|
| | **Maximum 37.48 Ma** | | **Maximum 41.2 Ma** | | **Maximum 52.4 Ma** | |
| **Node** | **AC** | **Independent** | **AC** | **Independent** | **AC** | **Independent** |
| C | 37.0 (36.4-37.5) | 37.0 (36.4-37.5) | 38.9 (36.5-41.2) | 38.9 (36.4-41.2) | 43.8 (36.1-51.8) | 44.4 (36.5-52.3) |
| M | 19.4 (17.8-21.3) | 19.2 (18.0-20.5) | 20.1 (17.9-22.5) | 20.1 (18.2-22.0) | 22.5 (18.1-27.3) | 23.0 (18.6-27.5) |
| 1 | 5.6 (4.9-6.3) | 4.3 (3.8-4.7) | 5.8 (5.0-6.6) | 4.5 (3.9-5.1) | 6.5 (5.1-8.0) | 5.1 (4.1-6.3) |
| 9 | 16.2 (14.8-17.9) | 16.2 (15.2-17.3) | 16.8 (15.0-18.9) | 17 (15.4-18.6) | 18.8 (15.1-22.9) | 19.4 (15.7-23.3) |
| 8 | 11.0 (10.0-12.1) | 11.1 (10.3-11.9) | 11.4 (10.1-12.9) | 11.7 (10.5-12.8) | 12.7 (10.2-15.5) | 13.3 (10.7-15.9) |
| 7 | 9.5 (8.6-10.5) | 9.6 (8.9-10.3) | 9.8 (8.7-11.1) | 10 (9.0-11.1) | 11.0 (8.8-13.4) | 11.4 (9.2-13.8) |
| 6 | 8.8 (8.0-9.8) | 9.0 (8.3-9.6) | 9.2 (8.1-10.3) | 9.4 (8.4-10.3) | 10.2 (8.2-12.5) | 10.7 (8.6-12.9) |
| 4 | 6.0 (5.4-6.7) | 6.1 (5.6-6.7) | 6.2 (5.5-7.1) | 6.4 (5.7-7.2) | 7.0 (5.6-8.6) | 7.3 (5.8-8.9) |
| 3 | 6.9 (6.2-7.7) | 6.9 (6.4-7.6) | 7.2 (6.5-8.2) | 7.3 (6.4-8.0) | 8.0 (6.4-9.9) | 8.3 (6.6-10.0) |

We constrained the minimum age of three nodes in our species tree: 36.4 Ma for the common ancestor of mysticetes and odontocetes (the root of the tree), based on *Mystacodon selenensis* (Lambert et al., 2017); 18.2 Ma for the common ancestor of Balaenidae, Balaenopteridae, and *Caperea*, based on *Morenocetus parvus* (Buono et al., 2017, Marx & Fordyce, 2015); and 8.37 Ma for the common ancestor of *Balaenoptera*, *Megaptera*, and *Eschrichtius*, based on *Archaeschrichtius ruggieroi* (Bisconti & Varola, 2006, Lirer et al., 2019).

Our *mcmctree* analysis was run six times using two different clock models – relaxed clock with autocorrelated rates and relaxed clock with independent rates – and three alternative maximum root ages: 52.4 Ma, based on the oldest known archaeocete *Himalayacetus subathuensis* (following (McGowen et al., 2020)); 41.2 Ma (base of the Priabonian), based on the absence of unequivocal crown cetaceans prior to the Late Eocene; and 37.48 Ma based on the upper bound of McGowen et al.'s (McGowen et al., 2020) 95% highest posterior density for the equivalent node (Table S2). Each *mcmctree* analysis was run for $10^6$ generations, sampling every 100 generations, following a 1% burn-in. To improve computational efficiency, analyses were run using an approximation of the maximum likelihood calculation (dos Reis & Yang, 2011). To ensure convergence of the MCMC, we ran duplicate chains for each analysis to compare the posterior estimates, and monitored the chains using Tracer v1.6 (Rambaut et al., 2018) to ensure effective sample sizes were >200 for each parameter.

*Pygmy right whale past demography*

We used the Pairwise Sequentially Markovian Coalescent (PSMC) v0.6.5 (Li & Durbin, 2011) to estimate past fluctuations in effective population size ($N_e$) of the pygmy right whale. The PSMC model identifies historical recombination events across a single diploid genome to infer the time to the most recent common ancestor (TMRCA) between genome segments. Assuming that pairwise sequence divergence is proportional to the timing of the coalescent events, parts of the genome with low heterozygosity indicate recent coalescence while regions of high heterozygosity correspond to more ancient coalescent events. Because the rate of coalescence is inversely proportional to $N_e$ (Li & Durbin, 2011), it can be used to estimate changes in $N_e$ through time.

We generated consensus sequences for all autosomes of the pygmy right whale genome from its bam file using the SAMtools *mpileup* (Li et al., 2009) command and the *vcf2fq* command of vcfutils.pl. We excluded sites with base and mapping quality <30 as well as sites with depth < ⅓ and > 2x the average coverage. We set N (the number of iterations) = 25, t (Tmax) = 15 and p (atomic time interval) = 64 (4 + 25*2 + 4 + 6, for each of which parameters are estimated with 28 free interval parameters) for the inference of TMRCA between each chromosome. We scaled population parameters assuming a generation time of 30 years and a substitution rate of $1.38e^{-08}$ substitutions/site/generation following (Árnason et al., 2018).

*Data deposition*

Trimmed reads for the pygmy right whale are available from the European Nucleotide Archive (PRJEB46892). For scripts, go to: https://github.com/ndussex/whale_genomics.git.
